# Changes in LXRα phosphorylation promote a novel diet-induced transcriptome that alters the transition from fatty liver to steatohepatitis

**DOI:** 10.1101/127779

**Authors:** Natalia Becares, Matthew C Gage, Lucia Martin-Gutierrez, Elina Shrestha, Rikah Louie, Benoit Pourcet, Oscar M Pello, Tu Vinh Luong, Saioa Goñi, Ning Liang, Cesar Pichardo, Hanne Røberg-Larsen, Vanessa Diaz, Knut R. Steffensen, Michael J. Garabedian, Krista Rombouts, Eckardt Treuter, Inés Pineda-Torra

## Abstract

Understanding the transition from fatty liver (steatosis) to inflammatory and fibrotic steatohepatitis, is key to define strategies that alter its progression. Here we show that, when challenged with a high fat-high cholesterol diet, mice carrying a mutation that abolishes phosphorylation at Ser196 (S196A) in the liver X receptor alpha (LXRα) exhibit reduced hepatic inflammation and fibrosis despite displaying enhanced steatosis. This is associated with a marked protection against cholesterol accumulation. Reduced steatohepatitis in S196A mice involves unique reprogramming of the liver transcriptome in response to the diet. Remarkably, impaired LXRα phosphorylation uncovers novel diet-specific/phosphorylation-sensitive genes, whose regulation does not simply mirror ligand-induced LXR activation. Regulation of these unique, dually responsive genes, is associated with the promotion of LXR and cofactor occupancy under a cholesterol-rich diet. Therefore, Ser196-LXRα phosphorylation acts as a novel nutritional sensor that triggers a unique diet-induced transcriptome, thereby modulating metabolic, inflammatory and fibrotic responses important in the transition to steatohepatitis.

## INTRODUCTION

Non-Alcoholic Fatty Liver Disease (NAFLD) involves conditions ranging from simple steatosis, steatosis accompanied by inflammation with or without fibrosis (steatohepatitis or NASH), progression to necrosis, cirrhosis and hepatocellular carcinoma promoting liver-related mortality (1). NAFLD is characterized by hepatic triglyceride and cholesterol accumulation without significant alcohol consumption. Steatosis alone is considered relatively benign, but its transition to NASH represents a key step into further liver damage, which without intervention can lead to organ transplantation. However, the mechanisms underlying this transition are poorly understood. NAFLD is strongly associated with obesity, insulin resistance and type 2 diabetes affecting 20-30% of the adult population in the West (1). Therapies such as insulin sensitizers, or lipid-lowering drugs are aimed at treating these associated conditions and only display limited efficacy. Indeed, effective NAFLD therapies are currently lacking and restricted to weight loss through lifestyle modifications (2). Therefore, identifying factors that modulate the transition to NASH is crucial for the development of novel treatments directly targeting NAFLD.

LXRα and LXRβ lipid-activated transcription factors heterodimerise with the Retinoid X Receptor (RXR) to control cholesterol and fatty acid homeostasis (3). In addition, LXRs act as modulators of inflammation and immunity (4) and show anti-inflammatory and anti-fibrotic activities in acute liver disease models (5,6). Besides ligand binding, LXR activity is modulated by post-translational modifications (7). We and others previously showed that LXRα is phosphorylated at Ser196 (Ser198 in the human homolog) (8–10) and that ligand-induced LXRα phosphorylation at this site alters its activity in a gene-specific manner (8,10). However, the physiological consequences of LXRα phosphorylation remain unknown. Here, we identify global LXRα phosphorylation at Ser196 as a nutritional sensor that critically impacts the transition to steatohepatitis in a dietary model of NAFLD.

## RESULTS

### LXRα-S196A mice exhibit enhanced steatosis

As we previously identified in macrophages (8,10), LXRα is phosphorylated at Ser196 (mouse) within a motif not present in LXRβ (Fig. S1A), in mouse (Fig. 1A) and human liver (Fig. S1B). To understand the impact of LXRα phosphorylation in response to a pathogenic diet, we generated a global knock-in mouse carrying a homozygous serine-to-alanine mutation at Ser196 (S196A) that impairs its phosphorylation (Fig. 1A and S1C-E). This has little effect on hepatic metabolism on a chow diet given that mutant mice have no apparent dysmorphic phenotype, display similar developmental growth to matching wild-type mice (WT) (not shown), and comparable hepatic lipids or other metabolic parameters on this diet (Table S1).

**Figure 1.**
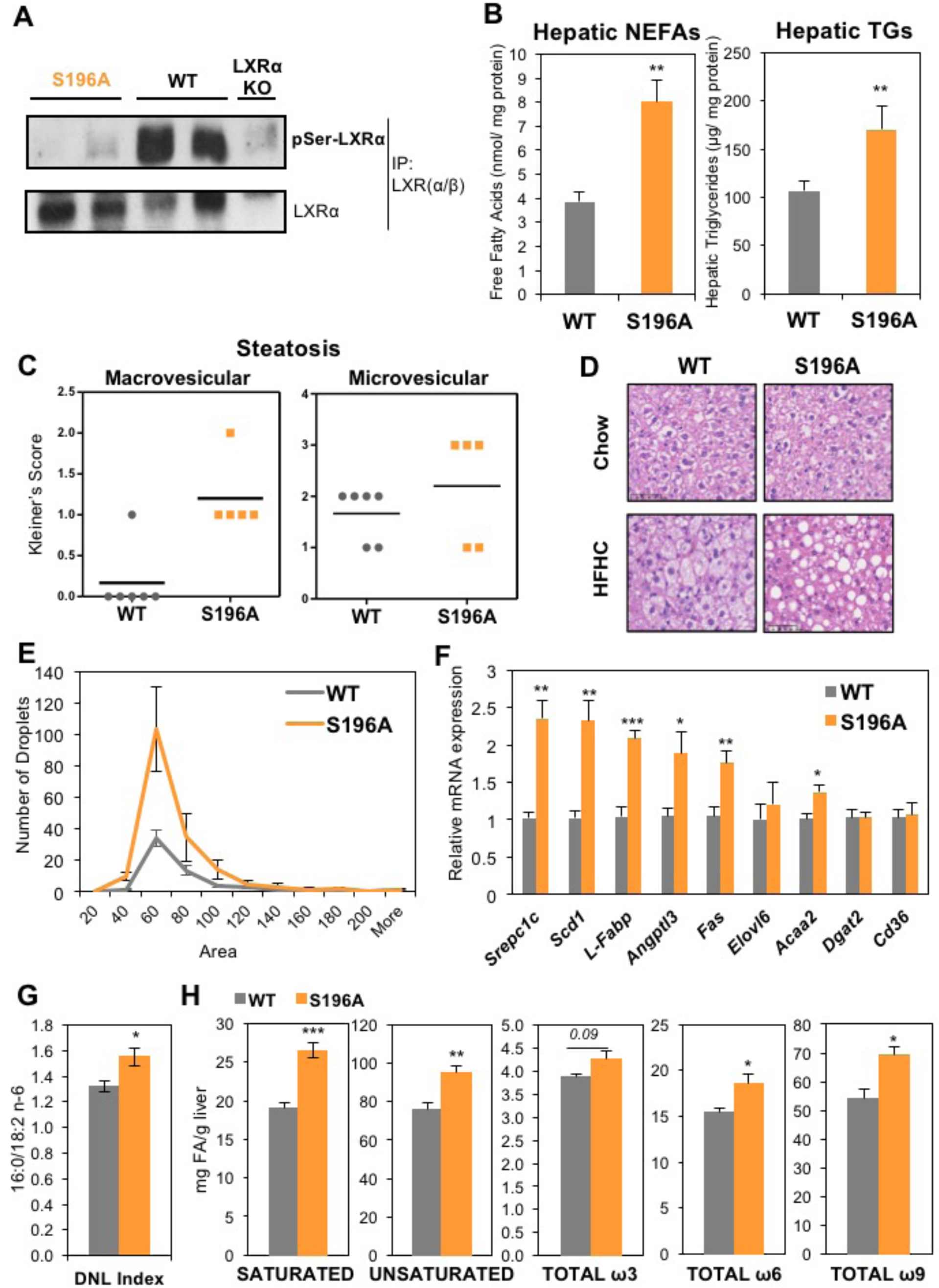
LXRα-S196A mice develop enhanced steatosis on a high cholesterol diet. (**a**) LXRα-Ser196 phosphorylation analysed by LXRα/β immunoprecipitation of liver homogenates and immunoblotting with a phospho-S196-LXRαspecific antibody. Global LXRα expression was assessed. (**b**) Hepatic non-esterified fatty acids (NEFAs) and triglycerides (TGs) in mice fed a HFHC diet (n=6/group) normalized to liver protein levels. (**c**) Kleiner’s scores for steatosis (0-3) of liver sections (n≥5/group). (**d**) Representative images of Haematoxylin and Eosin (H&E) stained liver sections from mice fed chow or HFHC diet. Scale bar at 50 μM. (**e**) Distribution of lipid droplets by area in H&E-stained liver sections (n=6/group). Area distribution was compared by chisquare test for trend (p= 0.0003). (**f**) Hepatic gene expression in mice fed a HFHC diet (n=6/group). Normalized data shown relative to WT, set as 1. (**g**) *De novo* lipogenesis (DNL) index measured as the ratio of 16:0 (Palmitate) and 18:2 n-6 (Linoleic) content in liver (n=6/group). (**h**) Hepatic fatty acid levels (n=6/group). Data are means ± SEM. *p<0.05, **p<0.005 or ***p<0.005 relative to WT.

Cholesterol metabolites are known LXR endogenous ligands (11) and diets with a high cholesterol content enhance LXR activity *in vivo* (12). Given that cholesterol induces LXRα phosphorylation (8), we hypothesised that LXRα phospho-mutant animals respond differently to a high fat-high cholesterol (HFHC) diet (13). Total body weight, plasma insulin and glucose levels were similar between S196A and WT mice fed a HFHC diet (Table S2). Consistent with LXRα role in hepatic steatosis (3), S196A mice displayed higher levels of hepatic, but not plasma, Non-Esterified Fatty Acids (NEFAs) and triglycerides than WT mice (Fig. 1B, S1F). Indeed, S196A mice showed enhanced micro and macrovesicular hepatic steatosis featuring more and larger lipid droplets (Fig. 1C-E) and enhanced expression of lipid droplet genes (Fig. S1G). Increased steatosis was associated with enhanced hepatic expression of the sterol response element binding protein 1 (*Srebp1c*) lipogenic transcription factor, and other LXR target genes involved in fatty acid synthesis (fatty acid synthase, *Fas*) (Fig. 1F). In contrast, other genes such as *Cd36,* involved in fatty acid uptake were not affected, recapitulating the gene selective effects of LXRα phosphorylation in macrophages we previously identified (8,10). Given that plasma NEFAs, triglycerides and insulin levels do not differ between genotypes (Fig. S1F and Table S2), increased hepatic fat accumulation in S196A mice likely results from enhanced *de novo* lipogenesis (Fig. 1G) as observed in other LXR models (14). S196A mice also showed an increase in the expression of stearoyl-CoA desaturase-1 (*Scd1*) (Fig. 1F), which catalyses the production of monounsaturated fatty acids. This led us to investigate whether changes in LXRα phosphorylation alter hepatic fatty acid composition, particularly since the types of fatty acids that accumulate in the liver during steatosis are thought to modulate the development of NAFL and its progression to NASH (15). Consistent with the changes in gene expression, S196A livers showed an increase in the total amount of saturated as well as unsaturated fatty acids, specifically ω9 and certain ω6 fatty acid species (Fig. 1h and Table S3). Altogether, these results demonstrate that LXRα phosphorylation deficiency at S196 induces hepatic steatosis and alters fatty acid composition in response to a HFHC diet.

### Impaired LXRα phosphorylation attenuates diet-induced hepatic inflammation and fibrosis

Diet-induced hepatic steatosis precedes inflammation and progression to fibrosis in experimental models (16). Strikingly, despite the increased steatosis, S196A mice displayed less inflammation (Fig. 2A) and significantly less collagen deposition than their WT counterparts (Fig. 2B). This was associated with a significant decrease in the expression of several pro-inflammatory and profibrotic mediators, such as *Oncostatin M* (*Osm*), *Chemokine (C-X-C motif) ligand 1* (*Cxcl1*) and *Osteopontin* (*Spp1*) and genes involved in collagen synthesis (*Col1a1* and *Tgfb2*) (Fig. 2C). Again, consistent with the reported gene specific effects on LXRα-modulated gene expression, only a subset of the genes analysed was affected by the S196A mutant (Fig. 2C and not shown). Notably, reduced levels of inflammatory and fibrotic genes in S196A mice were revealed mostly upon exposure to the cholesterol-rich diet while basal expression levels on chow were largely unaffected (Fig. 2D). This likely reflects a modulatory role for LXRα phosphorylation in diet-induced transcriptional responses.

**Figure 2.**
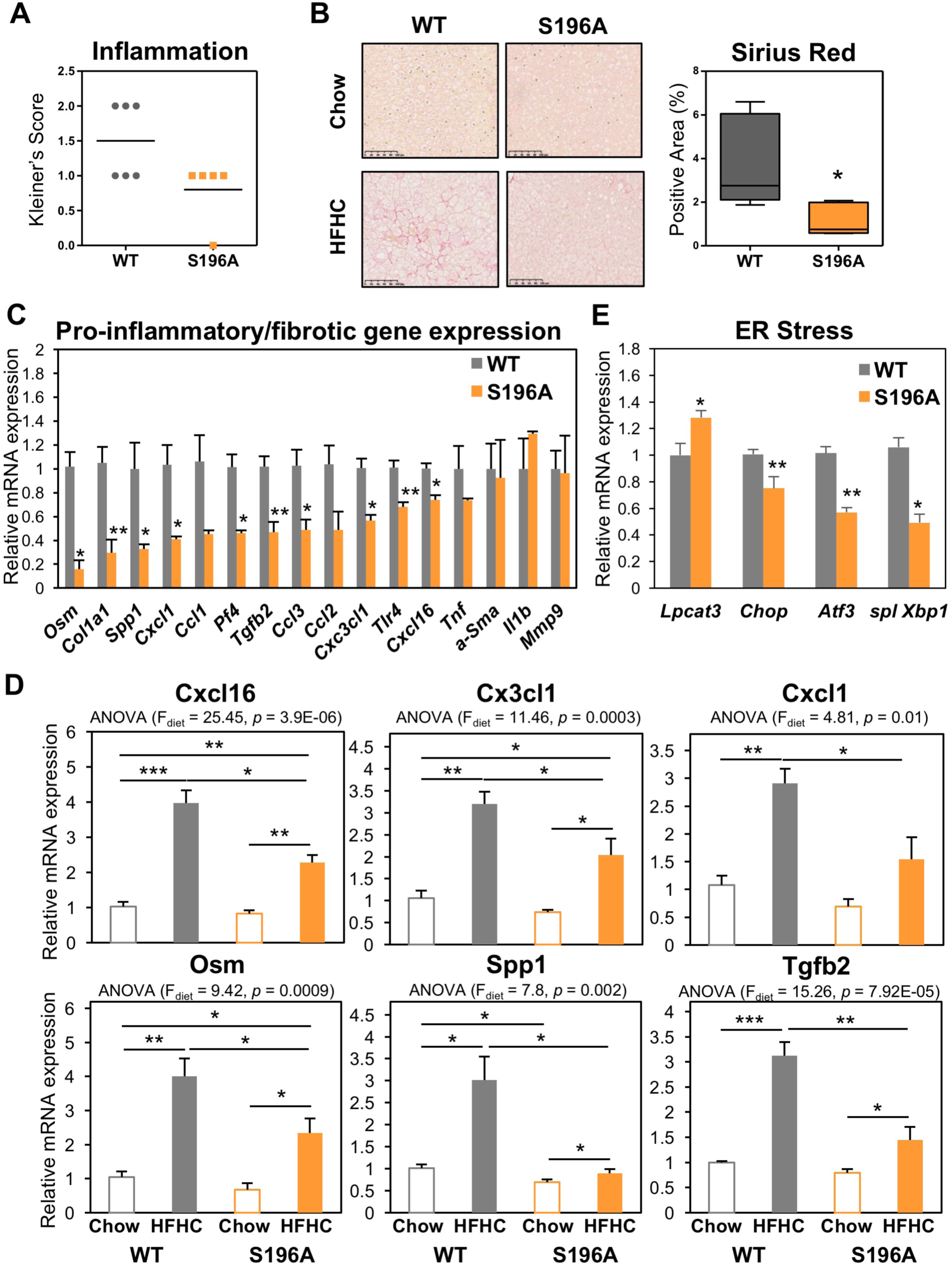
LXRα-S196A alleviates diet-induced hepatic inflammation and fibrosis. (**a**) Kleiner’s Scores for lobular inflammation (0-3) from liver sections of mice (n=6/group). (b) Representative images of Picrosirius Red stained liver sections (left). Scale bar at 100 μM. Quantification of stained areas by Image J (n=6/group) (right). Values are the average of positively-stained area. (c) Hepatic gene expression (n=6/group). Normalized data shown relative to WT. (d) Hepatic gene expression in mice fed chow (n=4/group) or HFHC diet (n=6/group). Normalized data shown relative to WT chow group. *p<0.05, **p<0.005, ***p< 0.0005, relative to WT chow. (e) Hepatic gene expression (n=6/group). Values shown normalized to cyclophilin and relative to WT. Data are means ± SEM. *p<0.05, **p<0.005 relative to WT.

Pathways implicated in the pathogenesis of lipid-induced liver damage, including apoptosis, lipid peroxidation and macrophage content, were similar between genotypes (Fig. S2A-C). In addition to these, prolonged adaptive endoplasmic reticulum (ER) stress is a known adaptive mechanism allowing cells to survive upon physiological changes requiring altered rates of protein folding. Notably, ER stress not only promotes steatosis but also modulates hepatic fibrosis (17).

Expression of factors involved in the activation of ER stress, such as the UPR target gene C/EBP homologous protein *(Chop*) and the Activating Transcription Factor (*Atf3*) or the spliced X-box-binding protein-1 (*Xbp-1*), was reduced in S196A mice (Fig. 2E), suggesting these animals could be protected from lipotoxicity through a reduction in ER stress. Overall, these findings demonstrate that blocking LXRα-phosphorylation at S196 attenuates lipid-induced hepatic inflammation and fibrosis despite the observed enhanced steatosis.

### LXRα phospho-mutant mice are protected from dietary cholesterol accumulation

Free cholesterol can act as an hepatotoxic agent (18) that induces collagen deposition in hepatic fibrosis (19). In striking contrast to WT mice, S196A mice challenged with a HFHC diet were protected from plasma and hepatic cholesterol accumulation (Fig. 3B). This was accompanied by a 20% reduction in liver weight in S196A livers compared to WT controls (Table S2). Thus, we next investigated expression of genes involved in cholesterol metabolism pathways that could be altered by the phospho-mutant LXRα, some of which are already well-characterised targets of LXRα (3). Reduced hepatic and plasma cholesterol levels were associated with decreased expression of cholesterol efflux transporter *Abcg1* (Fig. 3C), reflecting a dampened response to the cholesterol-rich diet (Fig. 3D). Notably, S196A mice showed a unique response to the HFHC diet regarding the upregulation of *Abcg5* (Fig. 3C,D), a transporter mediating hepatobiliary cholesterol secretion (20) and a well-characterised target of LXRα (3). No difference was observed in the levels of genes involved in cholesterol intestinal absorption and excretion (Fig. S2D), an important means by which LXR controls cholesterol homeostasis (3), nor in the expression of other nuclear receptors regulating lipid metabolism (Fig. S2E and not shown). Thus, the reduced cholesterol accumulation in S196A mice is likely due to increased hepatobiliary secretion of cholesterol. Moreover, in contrast to the strong repression of the cholesterogenic transcription factor *Srebp2* and its target gene *Ldlr* by dietary cholesterol seen in WT mice (Fig. 3E), these genes were largely unaffected by the diet in S196A mice, mirroring the unchanged hepatic cholesterol levels in these animals in response to diet (Fig. 3A). Overall, these differences in gene expression further reflect how the response to a cholesterol-rich diet differs between genotypes.

**Figure 3.**
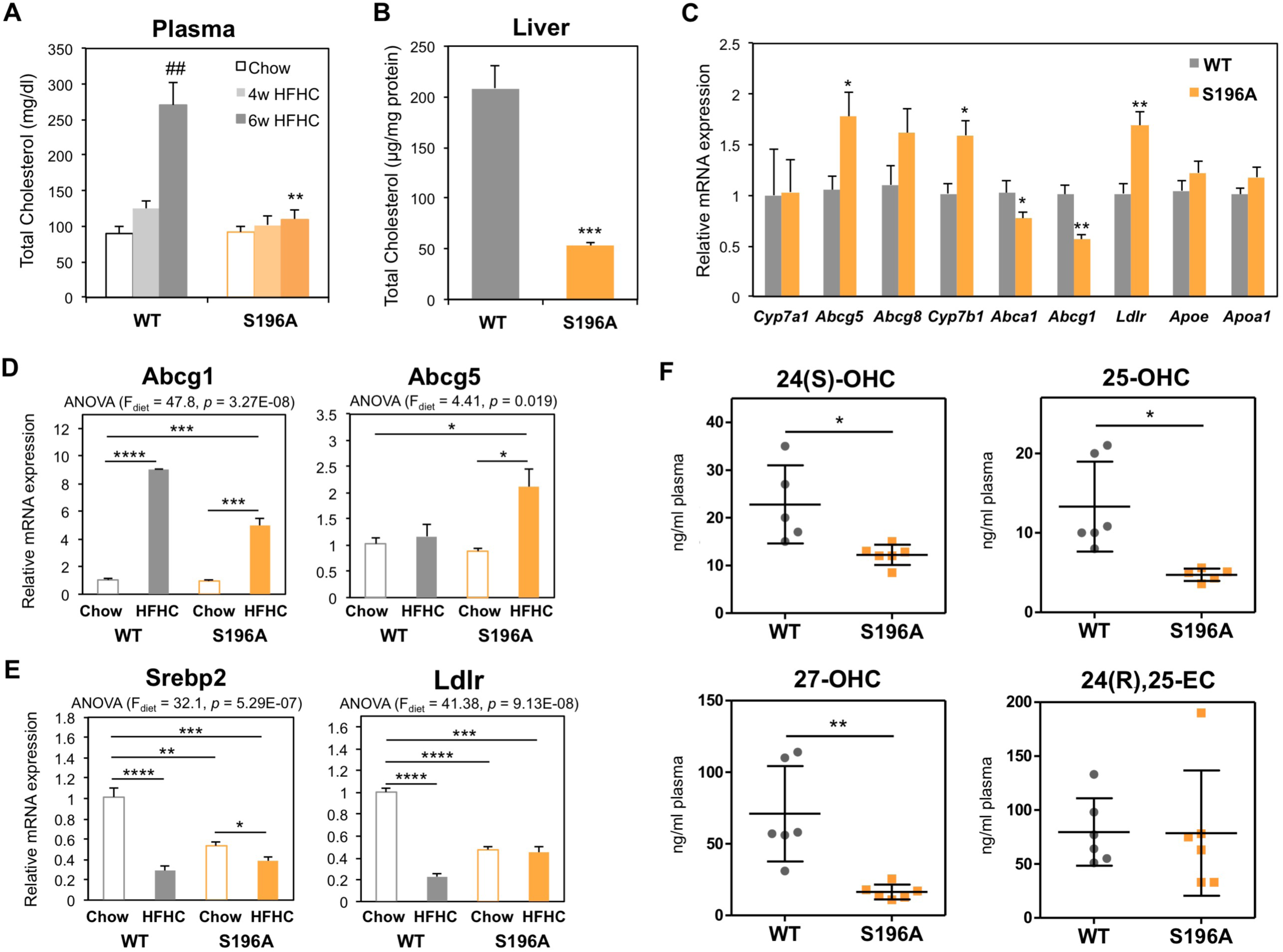
LXR phosphorylation deficient mice show reduced cholesterol levels in response to a HFHC diet. (**a**) Plasma total cholesterol levels in mice fed a chow (n=4/group) or a HFHC diet (n≥5/group). (**b**) Hepatic total cholesterol levels in mice fed a HFHC diet (n=6/group). Values shown are normalized to protein levels in tissue homogenates. (**c**) Hepatic gene expression in fed a HFHC diet (n=6/group). Normalized data shown relative to WT. (**d-e**) Hepatic gene expression in mice fed chow (n=4) or a HFHC diet (n=6). Normalized data shown relative to WT chow group. Significance of comparisons between HFHC WT and S196A genotypes is shown in c). (**f**) Quantification of free oxysterols in plasma of mice fed a HFHC diet (n=6/group). Data are means ± SEM. *p<0.05, **p<0.005, ***p<0.0005, ****p<0.00005 relative to WT. ##p<0.005, 4 vs 6 weeks.

Intracellular cholesterol accumulation activates the unfolded protein response pathway in the ER (21), inhibiting protein transport to the Golgi to re-establish ER function (22). One gene linking cholesterol metabolism and ER stress is *Tm7sf2*, which not only participates in cholesterol biosynthesis as a 3β-hydroxysterol Δ14-reductase but also acts as an ER sensor by triggering anti-inflammatory pathways (23). Consistent with a decreased hepatic inflammation (Fig. 2A), *Tm7sf2* expression was enhanced in S196A mice exposed to the diet while other genes involved in cholesterol biosynthesis were largely unaffected Fig. S2F). This suggests cholesterol modulation of ER stress responses, rather than cholesterol biosynthesis itself, is altered in S196A mice.

Because oxysterols (oxidised cholesterol derivatives some of which act as LXR ligands (11)) have been reported to be enhanced in NAFLD patients (24), we next examined the levels of these metabolites in both WT and S196A genotypes. LXRα phospho-mutant mice showed significantly reduced levels of most of the plasma oxysterols examined (Fig. 3F), consistent with the reduced signs of hepatic inflammation and fibrosis (Fig. 3C). These differences in oxysterol levels associated with induced *Cyp7b1* expression (Fig. 3C), an enzyme involved in oxysterol catabolism (25), while enzymes implicated in oxysterol synthesis remained unaffected (not shown). Overall, these findings suggest inhibition of LXRα phosphorylation acts as a novel molecular sensor of dietary cholesterol in the progression to steatohepatitis.

### LXRα-S196A reprograms hepatic gene expression and uncovers a novel diet-induced LXRα transcriptome

To better understand the disparity in diet-induced responses between genotypes and identify novel pathways sensitive to LXRα phosphorylation, we assessed transcriptomic differences by RNAseq analysis. Principal component analysis evidenced that transcriptomes of WT and S196A mice are substantially different, especially under a cholesterol-rich diet (Fig. 4A). Transcriptomic analysis revealed 668 genes whose hepatic expression is significantly altered in the mutant mice fed a HFHC diet (Fig. 4B,C and Fig. S3). Remarkably, there is minimal overlap between the genes modulated by LXRα phosphorylation at S196A in chow and HFHC diets, further reflecting on a genome-wide scale the distinct response exerted by S196A mice to the metabolic challenges posed by a cholesterol-rich diet. Pathway enrichment analysis confirmed our initial findings and showed induction of genes in lipid metabolism (Fig. 4D,E). Additional interrogation of our datasets revealed that in addition to increased expression of enzymes involved in fatty acid synthesis (*Srebf1, Fas,* Fig. 1F), S196A expression increased the hepatic levels of enzymes involved in fatty acid elongation (*Elovl3, Elovl5*) and fatty acid oxidation, with a trend towards increased levels of fatty acid desaturation enzyme *Fads1* (Fig. 4F). These changes likely contribute to the distinct hepatic fatty acid profile present in S196A mice (Fig. 1H). Interestingly, expression of most of these enzymes is severely repressed by the HFHC diet (Fig. S4a) further highlighting the modulatory role exerted by LXRα-S196. Also corroborating previous analyses, the phospho-mutant mice showed a robust decrease in the levels of wound healing and fibrotic mediators including several collagen genes and enzymes responsible for collagen stabilisation (Fig. 4D,G). Importantly, gene expression changes in response to a HFHC diet appeared to be substantially different between WT and S196A mice (Fig. S4B), further supporting that impaired phosphorylation of LXRα-S196 alters the susceptibility to diet-induced hepatic injury by inducing a distinct hepatic transcriptome. Moreover, expression of a subset of genes involved in extracellular matrix remodelling and tissue regeneration that distinguish between low-risk/mild and high-risk/severe NAFLD amongst pre-symptomatic patients (26) was remarkably different between genotypes (Fig. 4H). This suggests changes in LXRα phosphorylation could alter pre-clinical NAFLD progression. Importantly, most of these genes are involved in extracellular matrix remodelling and tissue regeneration, emphasizing a role for Ser196-LXRα phosphorylation in the regulation of these pathways.

**Figure 4.**
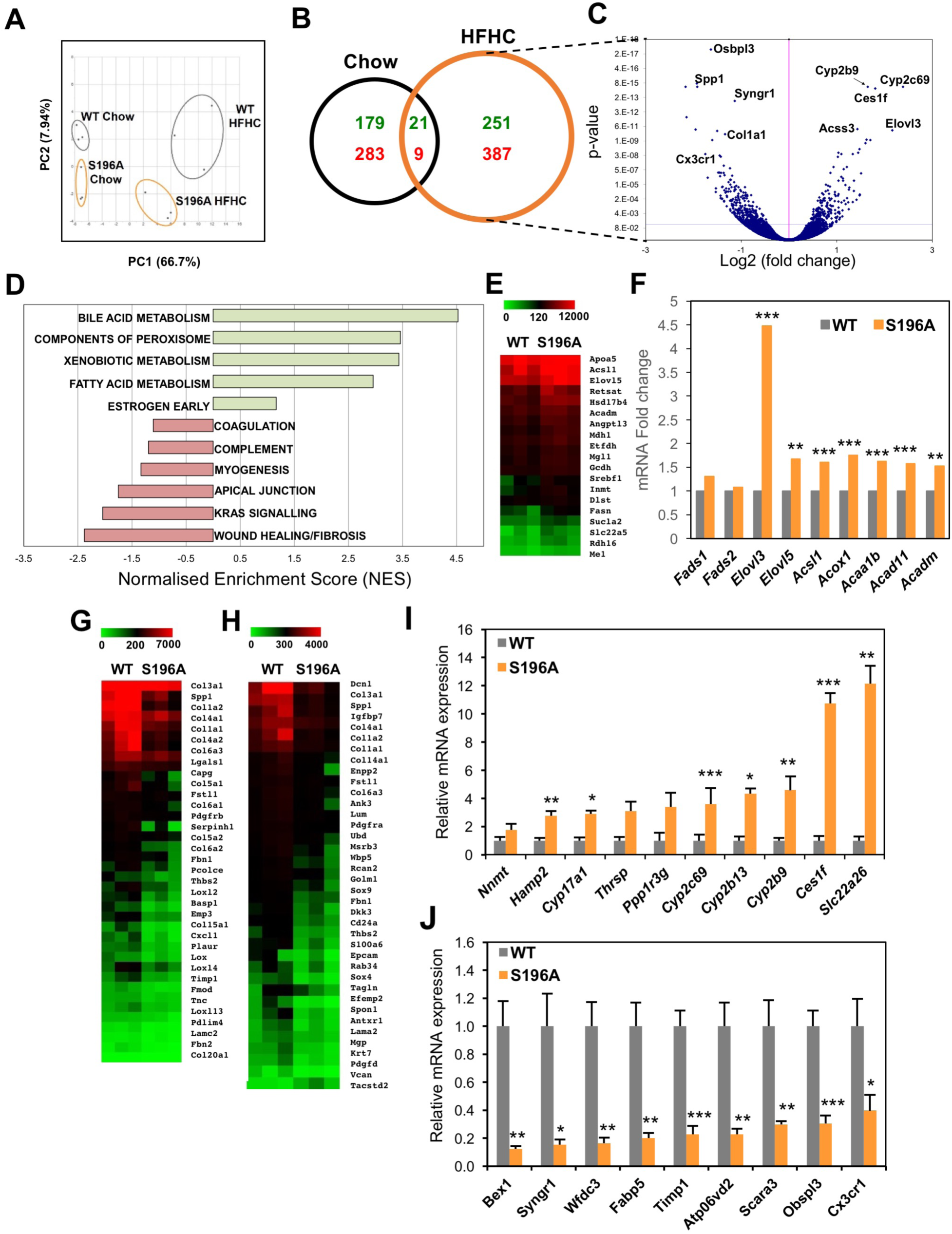
Changes in LXRα phosphorylation reprogram hepatic gene expression. (**a**) Principal Component (PC) Analysis plot showing RNAseq samples analysed by diet and genotype. (**b**) Venn diagram of genes regulated by LXRαS196A compared to LXRαWT (p<0.05) in the indicated diets. Numbers of upregulated and downregulated genes are depicted in green and red respectively. (**c**) Volcano plot of log_2_ ratio vs p-value of differentially expressed genes comparing S196A and WT livers exposed to a HFHC (n=3/group). Blue line indicates adjusted p-value threshold of 0.04 (Wald Test for logistic regression). (**d**) GSEA analysis showing enriched pathways in S196A livers with p<0.5 (100 permutations) derived from HALLMARK gene sets. (**e**) Heatmaps of hepatic RNAseq raw gene counts (n=3/genotype) for fatty acid and triglyceride metabolism, (**f**) Fold-change of hepatic RNAseq gene counts in S196A compared to WT mice fed a HFHC diet (n=3/genotype). Heatmaps of hepatic RNAseq gene counts (n=3/genotype) for (**g**) fibrosis and (**h**) human NAFLD signature genes. Hepatic gene expression by qPCR of top (**i**) upregulated and (**j**) downregulated genes from the RNAseq analysis on experimentally-independent WT and S196A livers (n=6/genotype). Normalized data shown relative to WT as mean ± SEM. *p<0.05, **p<0.005 or ***p<0.005 relative to WT.

Genes showing the strongest difference in expression between WT and S196A genotypes were confirmed in a separate set of mice (Fig. 4I,J). Notably, the majority of these genes are modulated by LXRα phosphorylation only in a cholesterol-rich environment (Fig. S4C,D) and have not been previously reported to be regulated by LXR. One such gene, *Ces1f*, is a member of the carboxylesterase 1 family that hydrolyses cholesterol esters and triglycerides and controls hepatic lipid mobilization (27,28). Despite previous studies showing *Ces1f* is not regulated by LXR ligands (29), we now clearly demonstrate *Ces1f* is highly sensitive to LXRα phosphorylation, preferentially in the context of a HFHC diet (Fig. 4I and S4C). Other Ces1 members are also differentially regulated by the LXRα phospho-mutant specifically in a cholesterol-rich setting (Fig S4E). Altogether, these data highlight the relevance of LXRα phosphorylation in modulating transcriptional responses to dietary cholesterol.

**Identification of novel LXR binding sites in dual LXRα phosphorylation/diet-sensitive genes** To better understand the differential gene regulation resulting from LXRαS196A expression, we next examined whether changes at LXRα phosphorylation at S196 affect LXR occupancy at selected genes in the context of a cholesterol-rich diet. *In silico* analysis of the *Ces1f* gene identified a DR4 sequence (AGGTCTatttAGTTCA), resembling the consensus LXRE (30), that was preferentially bound by LXR, but not RXR, in HFHC-fed S196A livers (Fig. 5A). This was associated with increased RNA Polymerase II (Pol II) and phosho-Ser2 Pol II (pSer-Pol II) occupancy indicating an enhanced transcriptional initiation and elongation respectively occurring at the *Ces1f* locus (Fig. 5D). A similar binding pattern was observed for a different DR4 sequence identified in *Cyp2c69* (GGGTCAagtgAGGTTA), another gene whose expression is enhanced in S196A mice (Fig. 4C,I) but not regulated by LXR ligands (31). In contrast, occupancy by both LXR and RXR to the well-established LXRE in *Srebp-1c* gene was induced (Fig. 5C), as was Pol II and pSer-Pol II (Fig. 5D). This shows impaired LXRαS196 phosphorylation allows the transcriptional activation of genes containing degenerated DR4 sequences without affecting RXR occupancy. The novel sequences identified here were revealed by the relative homology to reported LXREs in response to LXR ligands in the context of a chow diet (30), given these are the only ChIPseq LXR analyses available but bearing in mind that ligand-induced responses may not phenocopy binding patterns in response to a cholesterol-rich diet. Indeed, ChIPseq analysis in WT mouse livers treated with an LXR ligand on a chow diet shows very little to no presence of LXR at the *Ces1f* and *Cyp2c69* DR4 sequences identified (not shown), further supporting that both phosphorylation status of LXRα and diet environment are critical for the regulation of these genes.

**Figure 5.**
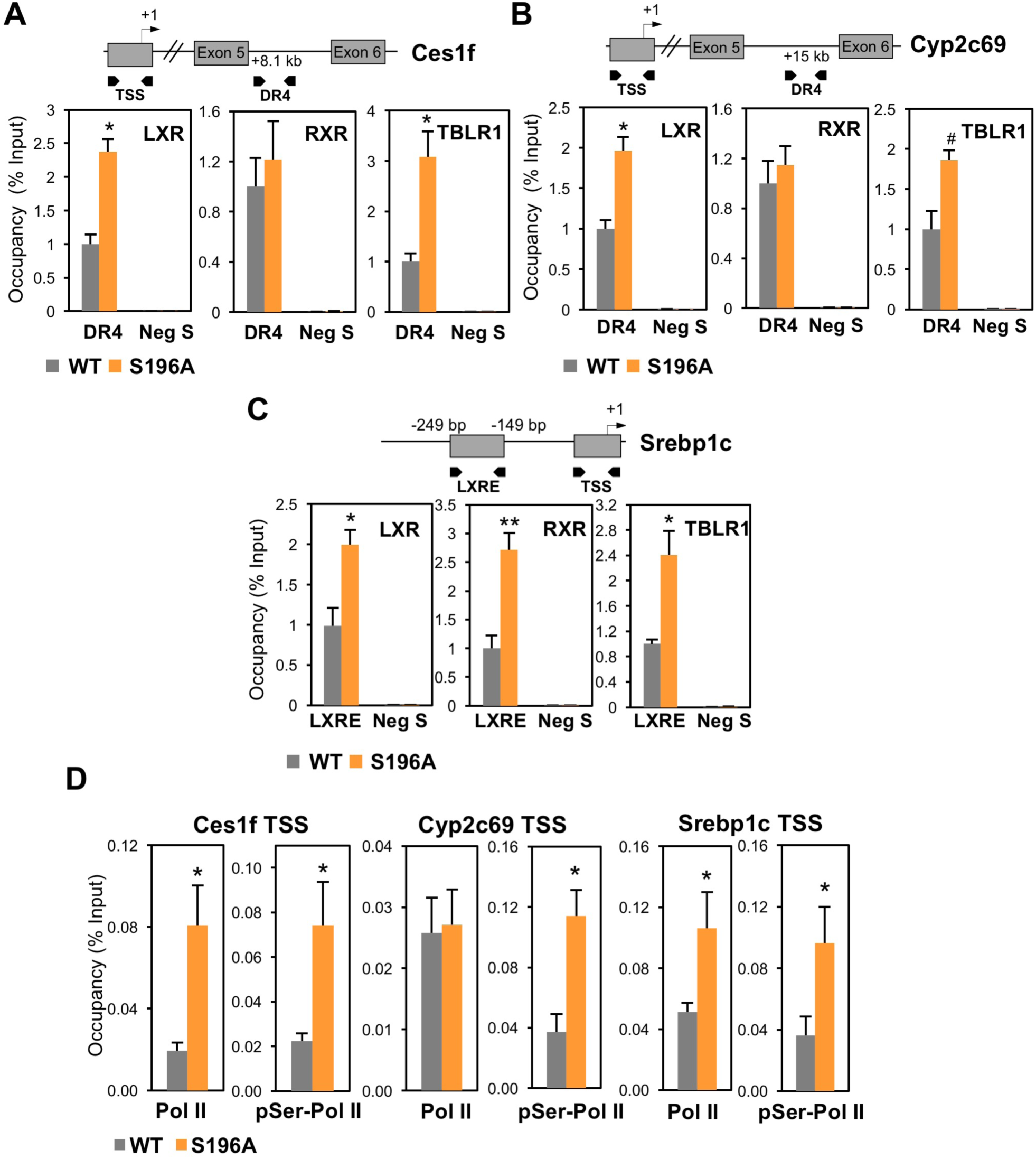
LXRα phosphorylation at S196 affects LXR and TBLR1 occupancy. (**a-c**) LXR, RXR and TBLR1 occupancy at Ces1f (**a**) and Cyp2c69 (**b**) identified DR4 sequences and Srebp-1c LXRE (**c**) or a region within in a gene desert (Neg S) in livers of mice fed HFHC for 6 weeks (n≥3/genotype). Results shown normalized to input and relative to WT. (**d**) RNA Pol II and pSer2-Pol II occupancy at Ces1f, Cyp2c69 and Srebp1c TSS in livers of WT and S196A mice fed a HFHC (n=3-6/genotype). Data represents mean ± SEM. #p=0.05, *p<0.05, **p<0.005 relative to WT.

**Figure 6.**
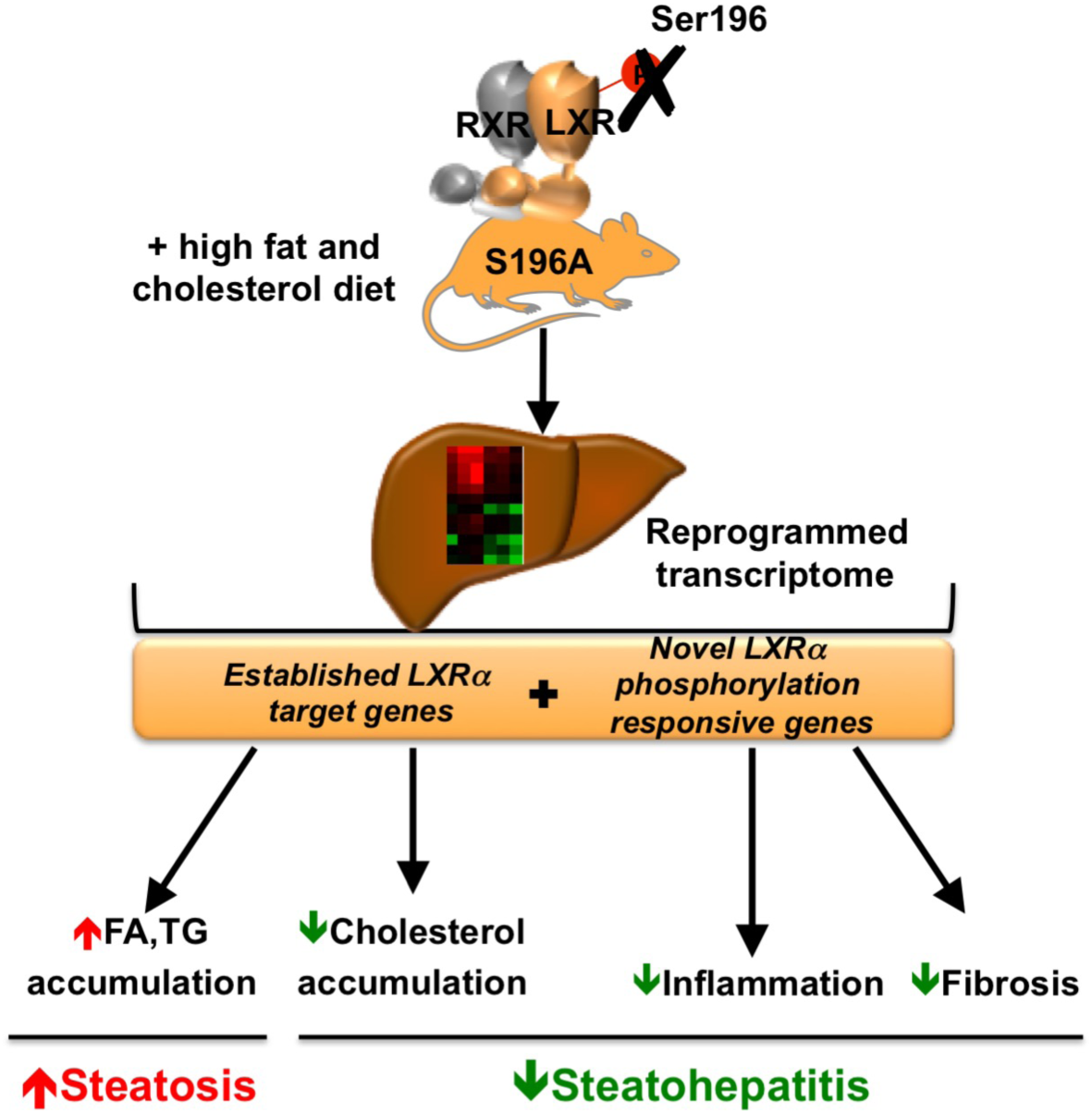
LXRα phosphorylation at S196 acts as a novel nutritional sensor and alters hepatic lipid metabolism. Impaired phosphorylation at LXRα-S196 reprograms the hepatic transcriptome in mice in response to a HFHC diet, leading to an increase in *de novo* lipogenesis and higher degree of steatosis, and counterintuitively, reduced hepatic inflammation and fibrosis (steatohepatitis).

Molecular modelling studies suggest that phosphorylation of LXRα at S198 (murine S196) induces a structural change in the hinge region of the receptor (8,10). Previously, we showed that upon LXR ligand activation, phosphorylation affects the transcriptional activity of LXRα by modulating the binding of the NCoR corepressor to phospho-sensitive genes (8). We were unable to detect differences in NCoR occupancy in mice exposed to the HFHC diet (not shown), suggesting responses to cholesterol *in vivo* involve other transcriptional players whose interaction with LXRα is sensitive to its phosphorylation status. Indeed, TBLR1, which participates in nuclear receptor cofactor exchange (32) and modulates LXR target gene expression in hepatic cells (33), was found to preferentially bind to LXRα-S196A (Fig. S5A,B). Consistently, TBLR1 occupancy was significantly enhanced in S196A livers exposed to the HFHC diet (Fig. 5A,B,C) suggesting this is an important component facilitating the transcription of these genes by the LXRα phospho-mutant in the context of a cholesterol rich diet. These data collectively indicate that disrupting LXRα phosphorylation at Ser196 affects diet-induced responses in liver and reveals LXR target genes likely through differential binding of LXR and TBLR1 to novel target sequences.

## DISCUSSION

The transition between relatively benign fatty liver steatosis to inflammatory and fibrotic steatohepatitis remains poorly understood. The role of LXRα in promoting fatty acid and triglyceride accumulation is well established (3) and has proven a major obstacle in the development of LXR ligands as therapeutics against metabolic and cardiovascular disorders (3). On the other hand, the hepatic anti-fibrotic and anti-inflammatory actions of LXRs in animal models of advanced fibrosis had shed light into additional pathways these receptors modulate in the progression to steatohepatitis (5,34). Based on the idea of reversing hepatic lipid accumulation, pharmacological antagonism of LXRs has been proposed as an effective therapy against NAFLD. For instance, a liver selective LXR inverse agonist SR9238 suppresses hepatic fatty acid synthesis and lipid accumulation leading to alleviated hepatic inflammation and fibrosis in an obese rodent model (35,36). However, it remained to be defined how LXR affects the transition to early fibrotic inflammatory stages of NAFLD in the context of a fatty liver, which is more clinically relevant. Previous studies focused their efforts at establishing changes in LXRα expression and reported induced levels of LXRα present in steatotic, inflammatory and fibrotic livers (37,38). These could, however, represent an adaptive or a maladaptive/pathogenic response to the ongoing cellular and molecular changes. Others studies however, have shown that LXRα is the only nuclear receptor whose expression is unaffected during progression to steatohepatitis (39). These contradictions highlight the need for further studies investigating how LXRs affect this chronic liver disease. We now propose that changes in LXRα phosphorylation play a crucial role in these transitional stages of the disease.

Regardless of changes in gene expression, changes in posttranslational modifications of the receptor could impact on the progression of the disease. Posttranslational modifications are a powerful means by which the activity and function of nuclear receptors can be altered. Despite the key importance of certain nuclear receptors in maintaining metabolic homeostasis, our understanding of how these modifications impact on metabolic diseases is scarce (7). Notably, the physiological consequences of LXRα phosphorylation, sumoylation and acetylation have only been studied *in vitro* or non-specifically in animal models by pharmacologically or genetically altering the enzymes enhancing or inhibiting these modifications (7). To directly address the impact of LXRα phosphorylation on NAFLD progression, we generated a novel mouse model harbouring an S196A mutation that disrupts LXRα phosphorylation at Ser196.

Here, we report for the first time that disrupting Ser196-LXRα phosphorylation affects hepatic diet induced responses by attenuating progression to steatohepatitis despite existing lipid accumulation. Importantly, LXRα phosphorylation at this residue dictates transcriptional responses to a HFHC diet that promotes early stages of NAFLD. Despite abundant triglyceride and NEFA accumulation, consistent with an increased *de novo* lipogenesis gene program (Fig. 1 & S1), S196A mice exhibit significantly less hepatic inflammation and fibrosis than WT animals (Fig. 2). This protective phenotype is associated with a dramatic reduction in hepatic and plasma cholesterol (Fig. 3) and a robust repression of numerous pro-inflammatory and pro-fibrotic mediators, including eleven collagen species, Lysyl oxidase (LOX) and lysyl oxidase-like proteins (LOXLs) responsible for collagen stabilisation through irreversible crosslinking (40,41) (Figs. 2 & 4). This is the first time Lysyl oxidase (LOX) and lysyl oxidase-like proteins (LOXLs) enzymes, which promote fibrosis progression and limit its reversibility (40), have been linked to LXRα.

Impaired LXRα phosphorylation uncovers novel diet-specific/phosphorylation-sensitive genes, *i.e.* genes responsive to changes in LXRα phosphorylation, mostly in the context of a cholesterol-rich diet (Fig. 4 & S4). Additionally, these genes have not been reported to be traditional ligand-modulated LXR targets suggesting the regulation of these identified genes does not simply phenocopy ligand-induced LXR activation. For instance, previous studies failed to show *Ces1f* regulation by LXR ligands (29) whereas we now demonstrate *Ces1f* is highly sensitive to LXRα phosphorylation in the context of early stages of steatohepatitis promoted by a HFHC diet (Fig. 4I & S4C). Ces1 has been recently shown to be protective from liver inflammation and injury (42) and its hepatic deficiency strongly increases susceptibility to cholesterol-driven liver injury (43). However, the specific contribution by the carboxylesterase 1 family member *Ces1f* in NAFLD progression has not been addressed. In addition to *Ces1f*, other Ces1 members (*Ces1b, Ces1c, Ces1d, Ces1e*) are differentially regulated by the LXRα phospho-mutant, most of which are only revealed to be sensitive to LXRα phosphorylation in a cholesterol-rich environment (Fig. S4E). Interestingly, the form previously shown to be induced by LXR ligands in liver, *Ces2c* (29), does not vary in S196A mice regardless of the diet (Fig S4E), emphasizing that LXRα phosphorylation-sensitive genes in response to diet are not necessarily responsive to LXR ligands and *vice versa*.

Impaired LXRα phosphorylation reveals unique LXRα phosphorylation/diet sensitive target genes associated with differential binding of LXR and TBLR1 to novel target sequences. Interestingly, our transcriptomic analysis showed that TBLR1 is neither regulated by changes in LXRα phosphorylation nor by exposure to the cholesterol-rich diet. TBLR1 activity itself is subject to regulation by posttranslational modifications. Perissi and colleagues demonstrated that TBLR1-S193 is a target of Protein Kinase Cδ-mediated phosphorylation, which happens in situ on the promoters of its regulated genes, and leads to the release and degradation of NCoR/SMRT (44). This further supports the idea that posttranslational modifications are used as a quick, reversible, targeted way to regulate the transcriptional machinery.

It is important to note that hepatic LXR binding at gene regulatory sites has never been explored in a cholesterol-rich diet setting before (30,45). Several recent studies have established that altered metabolic states promote epigenomic changes, through chromatin modifications both in animal models fed fat-rich diets and in obese and diabetic individuals(46–48). These modifications affect chromatin accessibility and are considered to act as a “metabolic imprint” that is able to alter metabolic disease risk, such as diabetes and NAFLD. Our findings suggest that the LXRα phosphorylation mutant takes advantage of these changes in the chromatin landscape to alter dietary transcriptional responses.

Overall, LXRα phosphorylation at Ser196 acts as a novel nutritional sensor that promotes a unique diet-induced transcriptome that modulates metabolic, inflammatory and fibrotic responses key in NAFLD progression. Understanding how this and other posttranslational modifications of LXRs are modulated and their impact on liver physiology could open alternative therapeutic avenues for NAFLD.

## METHODS

### Generation of the S196A transgenic animal models

The S196A floxed (S196A^fl/fl^) mouse line was generated by Ozgene Pty Ltd (Bentley WA, Australia). The genomic sequence for the murine LXRα (Nr1h3) gene was obtained from the Ensembl Mouse Genome Server (http://www.ensembl.org/Mus_musculus/), Ensembl gene ID: ENSMUSG00000002108. The mutant fragment, located on Exon 5, contains a serine-to-alanine mutation at Ser196 introduced by site-directed mutagenesis. The point-mutant exon was delivered into an intronic site inside the targeting vector, placed in opposite orientation and thus without coding capacity (Fig. S1A). The targeting construct was electroporated into the Bruce4 C57BL/6 ES cell line. Homologous recombinant ES cell clones were identified by Southern hybridization and injected into BALB/cJ blastocysts. Male chimeric mice were obtained and crossed to C57BL/6J females to establish heterozygous germline offsprings on a pure C57BL/6 background. The germline mice were crossed to a FLP Recombinase mouse line (49) to remove the FRT flanked selectable marker cassette (Flp’d mice). Flp’d mice were then crossed with a transgenic C57BL/6 mouse strain carrying a Cre recombinase under the PGK-1 promoter (50), resulting in the inversion and insertion of the lox-flanked mutated (loxP) vector exon 5 region in the sense orientation, and deletion of the wild-type (WT) sequence in most adult cell lineages (S196A mice) while WT matching controls carry the WT sequence in the sense orientation (Fig. S1D). Mice were genotyped by PCR analysis of ear biopsies (Fig S1D,E) using the Jumpstart Taq DNA Polymerase (Sigma Aldrich) and the following primers: wild-type (WT) forward 5’GGTGTCCCCAAGGGTGTCCT, reverse 5’ AAGCATGACCTGCACACAAG and mutant forward 5’ GGTGTCCCCAAGGGTGTCCG. Animals were housed together and maintained in a pathogen-free animal facility in a 12-h light-dark cycle. All procedures were carried under the UK’s Home Office Animals (Scientific Procedures) Act 1986.

### Diet studies and tissue collection

Ten-week old WT and S196A female mice were fed ad libitum a High Fat-High Cholesterol (HFHC) diet (17,2% Cocoa Butter, 2,8% Soybean Oil, 1,25% Cholesterol, 0,5% Sodium Cholate; AIN-76A/Clinton Diet #4, Test Diet Limited, UK) or a chow diet (18% Protein, 6.2% Fat, 0% Cholesterol; Harlan Laboratories) for 6 weeks. Mice were fasted overnight prior to sacrifice. Blood was collected by cardiac puncture and plasma was aliquoted and frozen at -80 °C. Tissue was dissected, weighted and frozen at -80 °C or placed in RNAlater (Sigma Aldrich).

### Plasma and hepatic lipid determination

Frozen livers (50 mg) were homogenized in 250 mM sucrose, 2 mM EDTA, 10 mM Tris buffer using ceramic beads in a Minilys Tissue Homogenizer (Bertin Corp.). Triglycerides and Cholesterol were extracted with Isopropanol or Chloroform:Methanol (1:1) solutions, respectively. Non Esterified Free Fatty Acids (NEFAs) were extracted by incubating liver homogenates with 1% Triton-100X and chloroform solution. Plasma and hepatic total cholesterol, triglyceride levels (Wako Diagnostics), and NEFAs (Abcam) were determined by colorimetric enzymatic assay kits as per the manufacturer’s recommendations. Hepatic lipid content was normalized to protein concentration.

### Oxysterol LC-MS analysis

Protein was precipitated from plasma with 480 pM of the appropriate internal standards (Avanti Polar Lipids, Alabaster, AL, USA). Sample clean-up was conducted off-line, using solid phase extraction (SPE, SilactSPE C18 100 mg, Teknolab, Ski, Norway) and dryed at 30 °C, re-dissolved in 2-propanol and treated as described (51). Samples and calibration solutions were analysed using an Ultimate 3000 UHPLC connected to an Advantage QqQ (both Thermo Fisher, Waltham, MA, USA) equipped with an Automatic filtration and filter back-flush SPE add-on, as described (52).

#### **Determination of hepatic fatty acid profiles**

Liver fatty acid content from WT and S196A mice fed a HFHC diet were assayed by gas liquid chromatography with flame ionization detection by AS Vitas (Oslo, Norway). Internal standard triheptadecanoin was added and fatty acids were methylated into methyl esters (FAMEs) with MeOH HCl and extracted with hexane. Analyses were performed on an Agilent 7890A GC and a 7683B automatic liquid sampler and flame ionization detection (Agilent Technologies, USA). Separations were obtained using a SP-2380 column. Fatty acid content was calculated based on the area percentage of peaks and response factors relative to 18:0. An external standard containing known amounts of relevant FAMEs (Supelco 37 component FAME Mix) was included in each run to correct for differences in fatty acid response factors. Results were normalized to protein content. The ratio of 16:0 to 18:2 n-6 was used to calculate a *de novo* lipogenesis index (53). The total saturated fatty acid content was calculated as the sum of 12:0, 14:0, 15:0, 16:0, 17:0, 18:0, 20:0 and 22:0. The total unsaturated fatty acid content was calculated as the sum of ω9 (16:1 c9, 18:1 c9, 20:1 n-9), ω6 (18:2 n-6, 18:3 n-6, 20:2 n-6, 20:3 n-6, 20:4 n-6) and ω3 (18:3 n-3, 20:5 n-3, 22:5 n-3, 22:6 n-3) fatty acids.

### RNA extraction and quantification

Total RNA from was extracted with TRIzol Reagent (Invitrogen). Sample concentration and purity was determined using a NanoDrop™ 1000 Spectrophotometer and cDNA was synthesized using the qScript cDNA Synthesis Kit (Quanta). Specific genes were amplified and quantified by quantitative PCR (qPCR), using the PerfeCTa SYBR Green FastMix (Quanta) on an MX3000p system (Agilent). Primer sequences are available upon request The relative amount of mRNAs was calculated using the comparative Ct method and normalized to the expression of cyclophilin (54). Mouse Cytokines & Chemokines RT2 Profiler PCR Arrays were performed per the manufacturer's instructions (Qiagen). Briefly, cDNA was synthesized using an RT^2^ HT first strand kit (Qiagen), and qPCR analysis was performed using RT2 SYBR Green ROX™ qPCR Mastermix (Qiagen). The relative amount of mRNAs was calculated using the comparative Ct method and normalized to an average of five housekeeping genes. The full list of genes analysed with these arrays can be found at Qiagen’s website.

### Protein isolation, immunoprecipitation, and immunoblotting

Single cell suspensions from livers were immunoprecipitated with antibodies that specifically recognise human (LXRα, ab41902 Abcam) or murine (LXRα/β) (55) receptors previously crosslinked to a column with Protein A/G Agarose following the manufacturer’s protocol (Pierce). Phospho-Ser196 specific rabbit polyclonal antibody (8), mouse α-LXRα monoclonal antibody (ab41902, Abcam), α-Hsp90 polyclonal (sc-7947, Santa Cruz) were used for immunoblotting. Anti-rabbit (PO448, Dako) or anti-mouse (NA931VS, GE Healthcare) horseradish-peroxidase-tagged antibodies were used for secondary binding and chemiluminescence (ECL 2 Western Blotting Substrate, Pierce) was used to visualise proteins.

For co-immunoprecipitation studies, HEK293T-LXRα, HEK-S198A or HEK 293T-Vo cells expressing FLAG-tagged receptors as in (8) were lysed and crude nuclear pellets were obtained. Supernatants containing nuclear proteins were incubated with FLAG antibody-conjugated agarose beads (Invitrogen). Bead-associated proteins associated were eluted in TBS and immunoblotted with α-TBLR1 (ab190796, Abcam) or α-LXRα (ab41902, Abcam) antibodies.

### Histopathological analysis

Formalin-fixed, paraffin-embedded mouse livers were cut and stained with hematoxylin and eosin (H&E) or Picrosirius Red (Abcam) dyes. Liver histology was blindly scored by an independent histopathologist based on three semiquantitative items: steatosis (0–3), lobular inflammation (0–3) and hepatocellular ballooning (0–2) (not shown) (56). Stained sections were scanned with NanoZoomer Digital slide scanner (Hamamatsu) and quantification of Picrosirius red-stained areas was performed using Image J on three independent areas per section. Data is represented as the average positively-stained percent of area of interest. Apoptosis was detected in liver tissue sections using a terminal deoxynucleotidyl transferase dUTP nick end-labeling (TUNEL) assay (R&D Systems). Sections were imaged using the Axio Imager.A1 Digital Microscope (Zeiss).

### Lipid droplet identification

Identification and quantification of lipid droplets were made with the help of Eli (Easy Lipids) v1.0, an in-house software developed between the Multiscale Cardiovascular Engineering (MUSE) and Dr Pineda-Torra’s groups at UCL. This software uses a method based on the Hough Transform (57) for the identification of the droplets estimating the centres and radii of each of them. A final report is generated with the dimensions of the droplets (i.e. diameter and area) including a histogram describing the frequency of lipid vacuoles within specified diameter ranges. A trial of Eli v1.0 is currently available upon request on the MUSE website at UCL (www.ucl.ac.uk/muse/software).

### Lipid peroxidation quantification

Thiobarbituric Acid Reactive Substances (TBARS) were measured in about 25 mg of frozen liver as per manufacturer’s instructions (Cayman Chemicals). Briefly, lipid peroxidation was quantified by the reaction of Malondialdehyde (MDA), a product of lipid peroxidation, with thiobarbituric acid (TBA) to form a colorimetric (532 nm) product, proportional to the MDA present. Levels of MDA were normalised to total protein levels, quantified by the Bradford Assay.

### LXRα proteomic analysis

HEK293T cells expressing vector only (Vo), FLAG-hLXRα or FLAG-hLXRα-S198A (58) were treated with 1 µM T0901317 for 8 hours. Purification of protein complexes, Multidimensional Protein Identification Technology, LTQ Mass Spectrometry and analysis was performed as described (58).

### Chromatin Immunoprecipitations

Fresh mouse livers were crosslinked with 2 mM disuccinimidyl glutarate (DSG) for 30 min, followed by 1% formaldehyde for 10 min at room temperature. The reaction was stopped with glycine at a final concentration of 0.125 M for 5 min. Single cell suspension were obtained by grinding liver pieces through a 70 μM cell strainer. Nuclei were isolated as described previously (59) and sonicated with the UCD-300 Bioruptor (Diagenode), to generate DNA-fragment sizes of 0.2–0.5 kb. The following antibodies were used for immunoprecipitations: RXRα (sc-553, Santa Cruz), Pol II (sc-9001, Santa Cruz), Pol II-S2P (ab5095, Abcam) and LXR (55). Following RNAse A (Fermentas) and proteinase K (Fermentas) treatment, immunoprecipitated DNA was purified using the QIAquick PCR purification kit (Qiagen) and analyzed by quantitative real-time PCR (primer sequences are listed in Table S3) and relative occupancies were normalized to input DNA (fold difference= 2 ^−Ct-sample-Ct-input^). To control for non-specific binding, a 82 base pair fragment in a gene desert in chromosome 6 (ActiveMotif) was used.

### RNA sequencing and analysis

Total RNA was extracted using TRIzol (Life technologies) and cDNA libraries were prepared using the Stranded mRNA-Seq Kit (Kapa Biosystems). Briefly, poly-A tailed RNA was purified using paramagnetic oligo-dT beads from 200 nanograms of total RNA, with a RNA Integrity Number above 7.5 as determined by the Agilent Bioanalyzer. The purified RNA was chemically fragmented and cDNA was synthesised using random primers (Kapa Biosystems). Adapter-ligated DNA library was amplified with 12 cycles of PCR and library fragment was estimated using the Agilent TapeStation 2200.Library concentration was determined using the Qubit DNA HS assay (Life Technologies). Libraries were sequenced on an Illumina NextSeq 500, NCS v2.1.2 (Illumina) with a 43bp paired end protocol. Basecalling was done using standard Illumina parameters (RTA 2.4.11). Sequencing and pipeline analysis was performed by UCL Genomics (London, UK). Reads were demulitplexed using Illumina’s bcl2fastq v2.17 and aligned using STAR v2.5.0b to the mouse GRCm38/mm10 reference sequence. Transcript abundance was estimated using Illumina's RnaReadCounter tool and differential expression analysis performed with DESeq2, which uses the Benjamin-Hochberg method for multiple testing correction. Pathway enrichment analysis was performed with the Gene Set Enrichment Analysis (GSEA) software’s pre-ranked module (60) and g:profiler (http://biit.cs.ut.ee/gprofiler/index.cgi). Top regulated genes were confirmed by qPCR on a separate set of liver samples from HFHC-fed mice. Heatmaps were created using raw gene count values with the MultiExperiment Viewer (MeV) software (61). Clustered heatmaps were created with Heatmapper Expression tool (<u>http://www1.heatmapper.ca/expression/</u>) and Venn diagrams using a BGE tool (<u>http://bioinformatics.psb.ugent.be/webtools/Venn/</u>).

### Statistical analysis

Data is presented as mean ± SEM. Differences were considered significant at *p* < 0.05 by a two-tailed Student t-test. For multiple comparisons, significance was assessed by single variance ANOVA followed by Student’s T-test. The *F*-statistic (df_between_=3, df_within_=15) and the *P* value for the significant main effect are shown.

### Data availability

Gene expression RNAseq data have been deposited at the NCBI Gene Expression Omnibus (GEO) accession numbers are GSE9665 (chow) and GSE95359 (HFHC). The authors declare that all data supporting the findings of this study are available within the article and its Supplementary Information files or are available from the corresponding author upon request.

## ACKNOWLEDGEMENTS

We are grateful to Prof Edward Fisher (NYU School of Medicine) and Dr James Thorne (University of Leeds) for insightful discussions. We also thank Dr Ruth Lovering (University College London) for her help with the bioinformatics analysis of the RNAseq datasets. This work was supported by a Medical Research Council New Investigator Grant G0801278 (IPT), British Heart Foundation Project Grant PG/13/10/30000 (IPT), UCL Grand Challenges PhD Studentship (NB, IPT), Swedish Research Council (VR2016-01743) (ET), Center for Innovative Medicine at Karolinska Institute (ET) and University of Oslo DIATECH@UiO initiative (HRL).

## AUTHOR CONTRIBUTIONS

N.B. performed most experiments, data analysis and prepared figures. M.G. helped to establish mouse colonies, performed experiments and data interpretation. L.M.G. and R.L. performed qPCR and analysed data. B.P. and O.M.P. helped establish mouse colonies. T.V.L scored liver sections. S.G., N.L. and E.T. provided unpublished ChIPseq data. C.P. and V.D. developed lipid droplet software. H.R-L. performed oxysterol analyses. K.R.S. and K.R. provided materials. M.J.G. provided materials and data and E.S. performed proteomic and immunoprecipitation analyses. K.R., M.J.G and E.T. helped with data interpretation. N.B. and I.P.T designed experiments, interpreted data and wrote the manuscript. I.P.T. conceived the study, secured funding and supervised all aspects of the work.

## References

1. EASL, EASD, EASO. EASL-EASD-EASO Clinical Practice Guidelines for the management of non-alcoholic fatty liver disease. J Hepatol. 2016;64(6):1388–402.

2. Thoma C, Day CP, Trenell MI. Lifestyle interventions for the treatment of non-alcoholic fatty liver disease in adults: A systematic review. J Hepatol. 2012;56(1):255–66.

3. Hong C, Tontonoz P. Liver X receptors in lipid metabolism: opportunities for drug discovery. Nat Rev Drug Discov [Internet]. Nature Publishing Group; 2014;13(6):433–44. http://www.ncbi.nlm.nih.gov/pubmed/24833295

4. Tall AR, Yvan-Charvet L. Cholesterol, inflammation and innate immunity. Nat Rev Immunol. Nature Research; 2015;15(2):104–16.

5. Beaven SW, Wroblewski K, Wang J, Hong C, Bensinger S, Tsukamoto H, et al. Liver X Receptor Signaling Is a Determinant of Stellate Cell Activation and Susceptibility to Fibrotic Liver Disease. Gastroenterology [Internet]. Elsevier Inc.; 2011;140(3):1052–62. http://dx.doi.org/10.1053Zj.gastro.2010.11.053

6. Hamilton JP, Koganti L, Muchenditsi A, Pendyala VS, Huso D, Hankin J, et al. Activation of liver X receptor/retinoid X receptor pathway ameliorates liver disease in Atp7B(-/-) (Wilson disease) mice. Hepatology [Internet]. 2016 Jun [cited 2016 Nov 30];63(6):1828–41. http://www.ncbi.nlm.nih.gov/pubmed/26679751

7. Becares N, Gage MC, Pineda-Torra I. Posttranslational Modifications of Lipid-Activated Nuclear Receptors: Focus on Metabolism. Endocrinology. 2016;158(2):213–25.

8. Torra IP, Ismaili N, Feig JE, Xu C-F, Cavasotto C, Pancratov R, et al. Phosphorylation of liver X receptor alpha selectively regulates target gene expression in macrophages. Mol Cell Biol. 2008;28(8):2626–36.

9. Chen M, Bradley MN, Beaven SW, Tontonoz P. Phosphorylation of the liver X receptors. FEBS Lett. 2006;580(20):4835–41.

10. Wu C, Hussein M, Shrestha E, Leone S, Aiyegbo MS, Lambert WM, et al. Modulation of macrophage gene expression via LXRa serine 198 phosphorylation. Mol Cell Biol [Internet]. 2015;35(11):2024–34. http://mcb.asm.org/content/35/11/2024.long

11. Janowski BA, Willy PJ, Devi TR, Falck JR, Mangelsdorf DJ. An oxysterol signalling pathway mediated by the nuclear receptor LXR alpha. [Internet]. Vol. 383, Nature. 1996. p. 728–31. http://www.ncbi.nlm.nih.gov/pubmed/8878485

12. Kalaany NY, Gauthier KC, Zavacki AM, Mammen PPA, Kitazume T, Peterson JA, et al. LXRs regulate the balance between fat storage and oxidation. Cell Metab. 2005;1(4):231–44.

13. Savard C, Tartaglione E V., Kuver R, Haigh WG, Farrell GC, Subramanian S, et al. Synergistic interaction of dietary cholesterol and dietary fat in inducing experimental steatohepatitis. Hepatology [Internet]. Wiley Subscription Services, Inc., A Wiley Company; 2013 Jan [cited 2017 Feb 6];57(1):81–92. http://doi.wiley.com/10.1002/hep.25789

14. Schultz JR, Tu H, Luk A, Repa JJ, Medina JC, Li L, et al. Role of LXRs in control of lipogenesis. Genes Dev. 2000;14(22):2831–8.

15. Peverill W, Powell LW, Skoien R. Evolving concepts in the pathogenesis of NASH: beyond steatosis and inflammation. Int J Mol Sci [Internet]. 2014 May 14 [cited 2017 Oct 11];15(5):8591–638. http://www.mdpi.com/1422-0067/15/5/8591/

16. Sanyal AJ. Mechanisms of Disease: pathogenesis of nonalcoholic fatty liver disease. Nat Clin Pract Gastroenterol Hepatol [Internet]. Nature Publishing Group; 2005 Jan [cited 2017 Feb 9];2(1):46–53. http://www.nature.com/doifinder/10.1038/ncpgasthep0084

17. Dara L, Ji C, Kaplowitz N. The contribution of endoplasmic reticulum stress to liver diseases. Hepatology [Internet]. NIH Public Access; 2011 May [cited 2017 Feb 14];53(5):1752–63. http://www.ncbi.nlm.nih.gov/pubmed/21384408

18. Mari M, Caballero F, Colell A, Morales A, Caballeria J, Fernandez A, et al. Mitochondrial free cholesterol loading sensitizes to TNF-and Fas-mediated steatohepatitis. Cell Metab. 2006;4(3):185–98.

19. Tomita K, Teratani T, Suzuki T, Shimizu M, Sato H, Narimatsu K, et al. Free cholesterol accumulation in hepatic stellate cells: Mechanism of liver fibrosis aggravation in nonalcoholic steatohepatitis in mice. Hepatology [Internet]. 2014 Jan [cited 2017 Feb 8];59(1):154–69. http://www.ncbi.nlm.nih.gov/pubmed/23832448

20. Wu JE, Basso F, Shamburek RD, Amar MJA, Vaisman B, Szakacs G, et al. Hepatic ABCG5 and ABCG8 overexpression increases hepatobiliary sterol transport but does not alter aortic atherosclerosis in transgenic mice. J Biol Chem. 2004;279(22):22913–25.

21. Devries-Seimon T, Li Y, Yao PM, Stone E, Wang Y, Davis RJ, et al. Cholesterol-induced macrophage apoptosis requires ER stress pathways and engagement of the type A scavenger receptor. J Cell Biol [Internet]. The Rockefeller University Press; 2005 Oct 10 [cited 2017 Feb 11];171(1):61–73. http://www.ncbi.nlm.nih.gov/pubmed/16203857

22. Kockx M, Dinnes DL, Huang K-Y, Sharpe LJ, Jessup W, Brown AJ, et al. Cholesterol accumulation inhibits ER to Golgi transport and protein secretion: studies of apolipoprotein E and VSVGt. Biochem J. 2012;447(1):51–60.

23. Bellezza I, Roberti R, Gatticchi L, Del Sordo R, Rambotti MG, Marchetti MC, et al. A Novel Role for Tm7sf2 Gene in Regulating TNFα Expression. Fessler MB, editor. PLoS One [Internet]. 2013 Jul 23 [cited 2017 Mar 7];8(7):e68017. Available from: http://www.ncbi.nlm.nih.gov/pubmed/23935851

24. Ikegami T, Hyogo H, Honda A, Miyazaki T, Tokushige K, Hashimoto E, et al. Increased serum liver X receptor ligand oxysterols in patients with non-alcoholic fatty liver disease. J Gastroenterol [Internet]. 2012 Nov 9 [cited 2017 Mar 6];47(11):1257–66. http://link.springer.com/10.1007/s00535-012-0585-0

25. Uppal H, Saini SPS, Moschetta A, Mu Y, Zhou J, Gong H, et al. Activation of LXRs prevents bile acid toxicity and cholestasis in female mice. Hepatology [Internet]. Wiley Subscription Services, Inc., A Wiley Company; 2007 Feb [cited 2017 Mar 2];45(2):422–32. http://doi.wiley.com/10.1002/hep.21494

26. Moylan CA, Pang H, Dellinger A, Suzuki A, Garrett ME, Guy CD, et al. Hepatic gene expression profiles differentiate presymptomatic patients with mild versus severe nonalcoholic fatty liver disease. Hepatology [Internet]. 2014 Feb [cited 2017 Mar 7];59(2):471–82. http://doi.wiley.com/10.1002/hep.26661

27. Zhao B, Natarajan R, Ghosh S. Human liver cholesteryl ester hydrolase: cloning, molecular characterization, and role in cellular cholesterol homeostasis. Physiol Genomics [Internet]. 2005 Nov 17 [cited 2017 Feb 3];23(3):304–10. http://www.ncbi.nlm.nih.gov/pubmed/16131527

28. Quiroga AD, Li L, Trötzmüller M, Nelson R, Proctor SD, Köfeler H, et al. Deficiency of carboxylesterase 1/esterase-x results in obesity, hepatic steatosis, and hyperlipidemia. Hepatology [Internet]. 2012 Dec [cited 2017 Feb 3];56(6):2188–98. http://www.ncbi.nlm.nih.gov/pubmed/22806626

29. Jones RD, Taylor AM, Tong EY, Repa JJ. Carboxylesterases are uniquely expressed among tissues and regulated by nuclear hormone receptors in the mouse. Drug Metab Dispos [Internet]. American Society for Pharmacology and Experimental Therapeutics; 2013 Jan [cited 2017 Feb 3];41(1):40–9. http://www.ncbi.nlm.nih.gov/pubmed/23011759

30. Boergesen M, Pedersen TA, Gross B, van Heeringen SJ, Hagenbeek D, Bindesboll C, et al. Genome-Wide Profiling of Liver X Receptor, Retinoid X Receptor, and Peroxisome Proliferator-Activated Receptor in Mouse Liver Reveals Extensive Sharing of Binding Sites. Mol Cell Biol [Internet]. 2012 Feb 15 [cited 2017 Feb 11];32(4):852–67. http://www.ncbi.nlm.nih.gov/pubmed/22158963

31. Lehrke M, Lebherz C, Millington SC, Guan H-P, Millar J, Rader DJ, et al. Diet-dependent cardiovascular lipid metabolism controlled by hepatic LXRα. Cell Metab [Internet]. 2005 May [cited 2017 May 1];1(5):297–308. http://www.ncbi.nlm.nih.gov/pubmed/16054077

32. Perissi V, Aggarwal A, Glass CK, Rose DW, Rosenfeld MG. A corepressor/coactivator exchange complex required for transcriptional activation by nuclear receptors and other regulated transcription factors. Cell [Internet]. 2004 Feb 20 [cited 2017 Mar 7];116(4):511–26. http://www.ncbi.nlm.nih.gov/pubmed/14980219

33. Jakobsson T, Venteclef N, Toresson G, Damdimopoulos AE, Ehrlund A, Lou X, et al. GPS2 Is Required for Cholesterol Efflux by Triggering Histone Demethylation, LXR Recruitment, and Coregulator Assembly at the ABCG1 Locus. Mol Cell. 2009;34(4):510–8.

34. Wouters K, van Bilsen M, van Gorp PJ, Bieghs V, Lütjohann D, Kerksiek A, et al. Intrahepatic cholesterol influences progression, inhibition and reversal of non-alcoholic steatohepatitis in hyperlipidemic mice. FEBS Lett [Internet]. Federation of European Biochemical Societies; 2010;584(5):1001–5. http://www.ncbi.nlm.nih.gov/pubmed/20114046

35. Griffett K, Solt L, El-Gendy B-D, Kamenecka T, Burris T. A liver-selective LXR inverse agonist that suppresses hepatic steatosis. ACS Chem Biol. 2013;8(3):559–67.

36. Griffett K, Welch RD, Flaveny CA, Kolar GR, Neuschwander-Tetri BA, Burris TP. The LXR inverse agonist SR9238 suppresses fibrosis in a model of non-alcoholic steatohepatitis. Mol Metab. Elsevier GmbH; 2015;4(4):353–7.

37. Ahn SB, Jang K, Jun DW, Lee BH, Shin KJ. Expression of liver X receptor correlates with intrahepatic inflammation and fibrosis in patients with nonalcoholic fatty liver disease. Dig Dis Sci [Internet]. 2014;59(12):2975–82. http://www.ncbi.nlm.nih.gov/pubmed/25102981

38. Lima-Cabello E, García-Mediavilla MV, Miquilena-Colina ME, Vargas-Castrillón J, Lozano-Rodríguez T, Fernández-Bermejo M, et al. Enhanced expression of pro-inflammatory mediators and liver X-receptor-regulated lipogenic genes in non-alcoholic fatty liver disease and hepatitis C. Clin Sci (Lond). 2011;120(6):239–50.

39. Aguilar-olivos NE, Carrillo-córdova D, Oria-hernández J, Sánchez-valle V, Ponciano-rodríguez G, Ramírez-jaramillo M. The nuclear receptor FXR, but not LXR, up-regulates bile acid transporter expression in non-alcoholic fatty liver disease. Ann Hepatol Off J Mex Assoc Hepatol. 2015;14(4):487–93.

40. Liu SB, Ikenaga N, Peng Z-W, Sverdlov DY, Greenstein A, Smith V, et al. Lysyl oxidase activity contributes to collagen stabilization during liver fibrosis progression and limits spontaneous fibrosis reversal in mice. FASEB J [Internet]. 2016 Apr 1 [cited 2017 Mar 7];30(4):1599–609. http://www.fasebj.org/cgi/doi/10.1096/fj.14-268425

41. Kanta J. Elastin in the Liver. Front Physiol [Internet]. 2016 Oct 25 [cited 2017 Mar 7];7:491. http://journal.frontiersin.org/article/10.3389/fphys.2016.00491/full

42. Xu J, Xu Y, Li Y, Jadhav K, You M, Yin L, et al. Carboxylesterase 1 Is Regulated by Hepatocyte Nuclear Factor 4α and Protects Against Alcohol-and MCD diet-induced Liver Injury. Sci Rep [Internet]. Nature Publishing Group; 2016 Apr 14 [cited 2017 Feb 3];6:24277. http://www.nature.com/articles/srep24277

43. Li J, Wang Y, Matye DJ, Chavan H, Krishnamurthy P, Li F, et al. Sortilin 1 Modulates Hepatic Cholesterol Lipotoxicity in Mice via Functional Interaction with Liver Carboxylesterase 1. J Biol Chem [Internet]. American Society for Biochemistry and Molecular Biology; 2017 Jan 6 [cited 2017 Mar 1];292(1):146–60. http://www.ncbi.nlm.nih.gov/pubmed/27881673

44. Perissi V, Scafoglio C, Zhang J, Ohgi KA, Rose DW, Glass CK, et al. TBL1 and TBLR1 Phosphorylation on Regulated Gene Promoters Overcomes Dual CtBP and NCoR/SMRT Transcriptional Repression Checkpoints. Mol Cell [Internet]. 2008 Mar 28 [cited 2017 Apr 20];29(6):755–66. http://www.ncbi.nlm.nih.gov/pubmed/18374649

45. Heinz S, Benner C, Spann N, Bertolino E, Lin YC, Laslo P, et al. Simple combinations of lineage-determining transcription factors prime cis-regulatory elements required for macrophage and B cell identities. Mol Cell [Internet]. 2010 May 28 [cited 2017 May 1];38(4):576–89. http://linkinghub.elsevier.com/retrieve/pii/S1097276510003667

46. Leung A, Parks BW, Du J, Trac C, Setten R, Chen Y, et al. Open chromatin profiling in mice livers reveals unique chromatin variations induced by high fat diet. J Biol Chem [Internet]. American Society for Biochemistry and Molecular Biology; 2014 Aug 22 [cited 2017 Jun 2];289(34):23557–67. http://www.ncbi.nlm.nih.gov/pubmed/25006255

47. Yuan W, Xia Y, Bell CG, Yet I, Ferreira T, Ward KJ, et al. An integrated epigenomic analysis for type 2 diabetes susceptibility loci in monozygotic twins. Nat Commun [Internet]. 2014 Dec 12 [cited 2017 Jun 2];5:5719. http://www.ncbi.nlm.nih.gov/pubmed/25502755

48. Leung A, Trac C, Du J, Natarajan R, Schones DE. Persistent Chromatin Modifications Induced by High Fat Diet. J Biol Chem [Internet]. American Society for Biochemistry and Molecular Biology; 2016 May 13 [cited 2017 Jun 2];291(20):10446–55. http://www.ncbi.nlm.nih.gov/pubmed/27006400

49. Takeuchi T, Nomura T, Tsujita M, Suzuki M, Fuse T, Mori H, et al. Flp recombinase transgenic mice of C57BL/6 strain for conditional gene targeting. Biochem Biophys Res Commun [Internet]. 2002 May 10 [cited 2017 Jan 23];293(3):953–7. http://www.ncbi.nlm.nih.gov/pubmed/12051751

50. Koentgen F, Suess G, Naf D. Engineering the Mouse Genome to Model Human Disease for Drug Discovery. In: Methods in molecular biology (Clifton, NJ). 2010. p. 55–77.

51. Roberg-Larsen H, Lund K, Vehus T, Solberg N, Vesterdal C, Misaghian D, et al. Highly automated nano-LC/MS-based approach for thousand cell-scale quantification of side chain-hydroxylated oxysterols. J Lipid Res [Internet]. 2014 Jul 1 [cited 2017 Mar 15];55(7):1531–6. http://www.jlr.org/cgi/doi/10.1194/jlr.D048801

52. Roberg-Larsen H, Lund K, Seterdal KE, Solheim S, Vehus T, Solberg N, et al. Mass spectrometric detection of 27-hydroxycholesterol in breast cancer exosomes. J Steroid Biochem Mol Biol [Internet]. 2016 Feb 10 [cited 2017 Mar 15]; http://linkinghub.elsevier.com/retrieve/pii/S0960076016300206

53. Chong MF-F, Hodson L, Bickerton AS, Roberts R, Neville M, Karpe F, et al. Parallel activation of de novo lipogenesis and stearoyl-CoA desaturase activity after 3 d of high-carbohydrate feeding. Am J Clin Nutr [Internet]. 2008 Apr [cited 2017 Oct 11];87(4):817–23. http://www.ncbi.nlm.nih.gov/pubmed/18400702

54. Pourcet B, Gage MC, León TE, Waddington KE, Pello OM, Steffensen KR, et al. The nuclear receptor LXR modulates interleukin-18 levels in macrophages through multiple mechanisms. Sci Rep [Internet]. 2016 May 6 [cited 2017 Feb 2];6:25481. http://www.nature.com/articles/srep25481

55. Pehkonen P, Welter-Stahl L, Diwo J, Ryynänen J, Wienecke-Baldacchino A, Heikkinen S, et al. Genome-wide landscape of liver X receptor chromatin binding and gene regulation in human macrophages. BMC Genomics [Internet]. 2012 Jan 31 [cited 2016 Dec 19];13(1):50. http://www.ncbi.nlm.nih.gov/pubmed/22292898

56. Liang W, Menke AL, Driessen A, Koek GH, Lindeman JH, Stoop R, et al. Establishment of a general NAFLD scoring system for rodent models and comparison to human liver pathology. PLoS One. 2014;9(12):1–17.

57. Duda RO, Hart PE. Use of the Hough transformation to detect lines and curves in pictures. Commun ACM [Internet]. ACM; 1972 Jan 1 [cited 2017 Feb 2];15(1):11—5. http://portal.acm.org/citation.cfm?doid=361237.361242

58. Shrestha E, Hussein MA, Savas JN, Ouimet M, Barrett TJ, Leone S, et al. Poly(ADP-ribose) Polymerase 1 Represses Liver X Receptor-mediated ABCA1 Expression and Cholesterol Efflux in Macrophages. J Biol Chem [Internet]. 2016 May 20 [cited 2017 Apr 5];291(21):11172–84. http://www.ncbi.nlm.nih.gov/pubmed/27026705

59. Fan R, Toubal A, Goñi S, Drareni K, Huang Z, Alzaid F, et al. Loss of the co-repressor GPS2 sensitizes macrophage activation upon metabolic stress induced by obesity and type 2 diabetes. Nat Med [Internet]. Nature Research; 2016 Jun 6 [cited 2017 Feb 2];22(7):780–91. http://www.nature.com/doifinder/10.1038/nm.4114

60. Subramanian A, Tamayo P, Mootha VK, Mukherjee S, Ebert BL, Gillette MA, et al. Gene set enrichment analysis: a knowledge-based approach for interpreting genome-wide expression profiles. Proc Natl Acad Sci U S A [Internet]. National Academy of Sciences; 2005 Oct 25 [cited 2017 Jan 20];102(43):15545–50. http://www.ncbi.nlm.nih.gov/pubmed/16199517

61. Howe EA, Sinha R, Schlauch D, Quackenbush J. RNA-Seq analysis in MeV. Bioinformatics [Internet]. Oxford University Press; 2011 Nov 15 [cited 2017 Mar 7];27(22):3209–10. http://www.ncbi.nlm.nih.gov/pubmed/2197642000

